# On the Depth of Deep Learning Models for Splice Site Identification

**DOI:** 10.1101/380667

**Authors:** Reem Elsousy, Nagarajan Kathiresan, Sabri Boughorbel

## Abstract

The success of deep learning has been shown in various fields including computer vision, speech recognition, natural language processing and bioinformatics. The advance of Deep Learning in Computer Vision has been an important source of inspiration for other research fields. The objective of this work is to adapt known deep learning models borrowed from computer vision such as VGGNet, Resnet and AlexNet for the classification of biological sequences. In particular, we are interested by the task of splice site identification based on raw DNA sequences. We focus on the role of model architecture depth on model training and classification performance.

We show that deep learning models outperform traditional classification methods (SVM, Random Forests, and Logistic Regression) for large training sets of raw DNA sequences. Three model families are analyzed in this work namely VGGNet, AlexNet and ResNet. Three depth levels are defined for each model family. The models are benchmarked using the following metrics: Area Under ROC curve (AUC), Number of model parameters, number of floating operations. Our extensive experimental evaluation show that shallow architectures have an overall better performance than deep models. We introduced a shallow version of ResNet, named S-ResNet. We show that it gives a good trade-off between model complexity and classification performance.

**Author summary:** Deep Learning has been widely applied to various fields in research and industry. It has been also succesfully applied to genomics and in particular to splice site identification. We are interested in the use of advanced neural networks borrowed from computer vision. We explored well-known models and their usability for the problem of splice site identification from raw sequences. Our extensive experimental analysis shows that shallow models outperform deep models. We introduce a new model called S-ResNet, which gives a good trade-off between computational complexity and classification accuracy.

## Introduction

The field of biology has witnessed, in the last decades, important advances thanks to major technological breakthroughs. Next Generation Sequencing (NGS) and other new tools have made fundamental changes in the understanding of numerous biological functions. The flood in biological data generated by NGS technology has motivated the use of data-driven approaches such as Deep Neural Networks to help understanding complex problems and extracting new biological knowledge. The ability of deep learning to cope with a variety of data formats has allowed handling biological sequences such as DNA, RNA or amino acid directly without a need for manual feature engineering. One of the challenges in bioinformatics is accurate identification of splice sites in DNA sequences. The discovery of splicing has elucidated the diversity of protein production and explained the increased coding potential of the genome. The DNA sequence is formed of alternating introns and exons, in the first stage, the DNA sequence transcribed into pre-mRNA, then, splicing process takes place by removing the non-coding sequences (introns) from the pre-mRNA to form mRNA sequence. However, a mutation could occur as a result of deletion or insertion of a number of nucleotides during the splicing process such as splicing in wrong site or removing exon region. This can result in a abnormal protein. Therefore the prediction of splice sites can provide valuable insights on the transcription process. The success of deep learning was originated from computer vision where typically a large number of training images are used. For example, the widely used ImageNet dataset gathers about 15 millions images categorized in about 1000 classes [1]. In order to learn from such a large amount of data deep learning architecture requires a large model capacity. Models with deep architecture have been proposed in the literature attaining 1000 layers [2–4]. However, this depth has raised a degradation problem presented in lower training accuracy when the number of layers extremely increased. Additionally, there are challenges in the training such models. ResNet models introduced residual blocks which connect with either identity mapping or projection shortcut [3]. ResNet structure helps to ease train the deep architecture and obtaining a good accuracy even with large number of layers.

Deep learning model architecture are designed based on the learning task, number of the parameters and size of the dataset. Well-known deep learning models, e.g., ResNet and VGGNet, from computer vision [2] have been reused to build advanced systems for text processing such as Very Deep Convolution Network (VDCNN) c [5] operating at character level directly. Text modeling and sentence classification have been also tackled with a small number of convolution layers such as one layer, two layers and six layers [6–8]. In recent work, the trend has gone toward evaluating the impact of the depth for text classification [7] and enhance the available techniques like generalizing the max-pooling operation and replace it with K-max pooling [7] resulting in Dynamic Convolution Neural Networks (DCNN) for semantic modeling of sentences.

Deep Learning has been also applied to splice site identification and has shown good performance compared to other machine learning approaches [9–11]. However most of existing deep learning methods are limited to simple models and no advanced architectures have been evaluated or proposed.

In this paper, we explore well-known deep learning architectures that were initially introduced for computer vision tasks and adapt them for the prediction of splice sites. The findings of this work can be useful for other applications in Bioinformatics. We selected three known deep learning models: VGGNet, AlexNet and ResNet. This choice is representative among the large number of successful models in the literature. Their architectures have different number of convolution layers between 5-6 in AlexNet 8-16 in VGGNet and 18-34 in ResNet. This paper contributes, in the topic of classification of splice sites, on several folds: 1) adaption of computer vision models such as VGGNet, ResNet and AlexNet for sequence data classification 2) Investigation of depth role on classification performance. 3) Proposal of a shallow architecture with residual shortcut (S-ResNet) based on ResNet leading to a best trade-off between model complexity and classification accuracy. 4) Benchmark of S-ResNet with traditional machine learning methods.

## Methods

Previous work in deep learning relied mainly on shallow architectures to classify DNA sequences and genomic data. In this work, we explored deeper architectures for splice site identification. We selected three well-known architecture families borrowed from computer vision to classify the splice sites. These three deep learning models are AlexNet [12], VGGNet [2], and ResNet [3]. Their architectures are described in the next sub-section. We kept the models as close as possible to original work. We mainly adapted them to our problem by using 1D convolutions layers and different number of filters. More details on the architectures can be also found in the original work previously cited. For each architecture family, we defined three models with shallow, intermediate and deep version of the architecture model. The depth is defined by the number of convolution layers in the model. We mention that deep and intermediate models have been used in the original work but the shallow versions are our adaptation inspired by the original models. Table 1 shows each original model with its intermediate and shallow version.

**Table 1.**
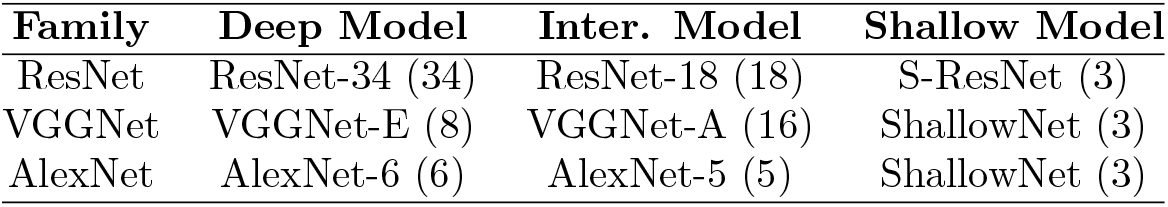
Three levels of depths for model families VGGNet, AlexNet and ResNet. These three levels are: Shallow, intermediate (Inter.) and Deep model. The number between parentheses depicts the number of convolution layers used.

| Family | Deep Model | Inter. Model | Shallow Model |
| --- | --- | --- | --- |
| ResNet | ResNet-34 (34) | ResNet-18 (18) | S-ResNet (3) |
| VGGNet | VGGNet-E (8) | VGGNet-A (16) | ShallowNet (3) |
| AlexNet | AlexNet-6 (6) | AlexNet-5 (5) | ShallowNet (3) |

### Experimental Setup

The Experimental evaluation is divided into four objectives which are detailed in the following points (1) Comparison of the proposed model S-ResNet with respect to traditional machine learning techniques such as Support Vector Machine (SVM), Logistic Regression (LG) and Random Forest (RF) (2) Analysis of deep learning model behaviour in the case of lengthy training setup. This aims mainly at exploring the behavior of ResNet models which are known to improve with long training. In these experiments, the number of epochs was fixed to 300. The experiments were repeated 5 times based on random splits of training, validation and test sets to estimate performance mean and standard deviation. This number was limited to reduce the computational time (3) The models are trained with early stopping based on monitoring the validation loss with patience set to 10 epochs. These evaluations were repeated 10 times (4) In this experiment we explored very long training for ResNet model. ResNet-18 was trained for 1000 epochs. The experimental results were analyzed based on different metrics measured during the experiments. These metrics are: accuracy, ROC Area Under Curve (ROC-AUC), number of model parameters, number of computation operations, sensitivity, and specificity. Both accuracy and AUC should be read as a percentage in which the value closer to 100 indicates better performance.

### Hyper-parameters Choice

The hyper-parameters were set, manually based on the validation set, after few trials to find the best combination. The choices were unified for all the models as much as possible. This is to remove external factors that could affect the evaluation and to keep mainly the model depth for comparison. The dataset was split into training and validation set with the percentage of 0.4 and 0.2 respectively to form 0.6 from the original dataset. The rest was used as test set. All models were trained with he_normal initialization [13] with a batch size of 250. ReLU was used as activation function and the last layer was fully connected with two nodes and softmax as activation function. Stochastic Gradient Descend (SGD) [14] was used to optimize the cross-entropy loss function with a learning rate of 0.01. The number of filters in the first convolution layer is chosen to be 8. This number is doubled for each next block. We define a block by any number of convolution layers between two max-pooling layers. Dropout with fraction of 0.5 was used as a regularizer with most of the architectures as detailed in the architecture sub-section.

### Splice Site Dataset

All the experiments were trained and tested using Splice Site Recognition (SSR) dataset. The task is to locate the boundaries between coding sequence (exons) and the non-coding (introns). To this end, splicing processing takes place in DNA sequence to strip the intron from pre-mRNA that is produced from transcription the DNA sequence. To ensure an accurate splicing, each exon is surrounded by splice-doner and splice-acceptor to determine the beginning and the end of each exon. In this work was splice sites are predicted given annotated DNA sequences with acceptor site and others not. The original dataset contained 159,771 true acceptor splice site sequences and 14,868,555 decoys (non-acceptor) and all the sequences with a length of 141 base pair. The same dataset has been previously used and is publicly available [15]. In order to remove the class imbalance problem, we used all the acceptor sequences and selected randomly about the same number of non-acceptor sequences (150,000). Thus, the total number is 309,771 DNA sequence samples. For benchmark with traditional methods, seven sub-datasets were selected randomly with different sizes (1000; 3000; 10,000; 30,000; 100,000; 200,000; 300,000). This aims at evaluating the effect of training dataset size on performance. For the other evaluations 300,000 examples were selected randomly from the full dataset. The data was split into three sets of training, validation, and test datasets respectively of size 0.4, 0.2, 0.4.

### Models Architecture

- **AlexNet**: AlexNet consists of two convolution layers each one followed by max-pooling and then either 3 or 4 convolutions layers called AlexNet_5 and AlexNet_6 respectively [12]. In this work the model suffix number indicates the total number of convolution layers. The model is ended with three fully connected layers. Rectified Linear Unit (ReLU) activation function has been used with all the layers. The output of ReLU has been normalized after the first and second convolution layers only. Overlap pooling approach was adopted to use a pooling with size z=3 and stride s=2 (siz) rather than local pooling (s=z). That is usually used to down sample the network. Three max-pooling layers have been used in the model, after the first, second and fifth convolution layers. The detailed model architecture is provided in Figure S.2 in the Supplementary material. In order to prevent over-fitting, dropout is applied after the first and second fully connected layers. The number of convolution filters starts with 8 filters (unless if we found more than this number can result in better performance) and then doubled each next convolution layer.
- **VGGNet**: VGGNet has a a slightly different configuration than AlexNet [2]. The number of convolution filters started with 8 and was doubled in each next block. In addition, stride of 1 was used. This is justified by giving importance individually to each nucleotide in the sequence. The original VGGNet in has many versions with different numbers of convolution layers, as of 8, 10, 13 or 16 layers separated with 5 max-pooling layers in between with the pooling size z=2 and stride s=2 followed with three fully connected layers [2]. In this work, the shallow and deep model were evaluated only with 8 and 16 convolutions (named VGGNet_A and VGGNet_E following the naming used in the original work). The convolution layers between two max-pooling layers are considered as a block and has the same number of filters. This number is doubled in the following block. In this work, the number of filters started with 8 with the same z=3 and s=1 in all the layers. Dropout function with ratio=0.5 for regularization the training after the first two fully connected layers.
- **ResNet**: The novelty in ResNet [3] is the introduction of a residual shortcut between blocks. There are two types of shortcuts: the fist one is known as identity shortcut where two connected layers have the same dimension. Thus, no extra dimensionality mapping are required. The second type is when the shortcut connects two layers with different dimensions. In this case, identity shortcut is still applicable but with extra zero padding. Alternatively, projection shortcut can be used. While projection shortcut equalizes the shape of the connected layers, it adds extra trainable weights to the model. In the original work, and also in this one, architectures with 18 convolutions and 34 have alternated identity and projection shortcuts as in Figure S.2. Batch normalization between the layers was used to avoid gradient vanishing and to maintain the forward signal with non-zero variances. The network is regularized by a dropout of 0.5 before every projection shortcut connection. The layers started with 16 filters with the z=3 and s=1. Similarly to the previous model families, all convolution layers have the same number of filters within a block and the number of filters is double in next blocks.
- **ShallowNet**: AlexNet and VGGNet have very similar architecture for shallow networks. Therefore we defined ShallowNet as a common representation for shallow version of both AlexNet and VGGNet. ShallowNet has three convolution layers, the first one followed by max-pooling then two convolution layers followed by the second max-pooling (see Figure S.2). Three fully-connected layers were used: The first and second one followed by a dropout layer with fraction 0.5. As in AlexNet, the output of the first convolution is activated by ReLU and then normalized. Conv.1 has 8 filters of the z=7 and s=1 and doubled in conv.2 and conv.3 with z=5 and s=1. Two overlapped max-pooling was used after conv.1 and conv.3 with z=3 and s=2.
- **DeepBind**: DeepBind is a well-known model for the prediction of sequence specificity of DNA and RNA binding protein [9]. We used it as an example for the simplest and shallowest architecture of DNA sequence classification. The model architecture is formed by one convolution layer with a ReLU activation and 8 filters with z=7 and s=1 of followed by a max-pooling with z=3 and s=2. Two fully-connected layers was used with a dropout of 0.5.
- **S-ResNet**: We propose a new model (S-ResNet) based on a shallow version of ResNet. It is different for ResNet by the use of a shortcut connection at each convolution layer whereas in ResNet the shortcut is placed after a block consist of two convolution layers. S-ResNet model architecture is depicted in Figure 1. It consists of three convolution layers and two overlap max-pooling with z=3 and s=2 after conv.1 and conv.3. All outputs of the convolution layers were normalized before the activation with a ReLU function. Two fully-connected layers were used without dropout. Three shortcuts were used two projection and one identity shortcuts. The projection shortcut was used to equalize the layers dimension between connected layers by the shortcut. The difference between S-ResNet and ResNet is the use of block with one convolution instead of two. In addition, shortcut are introduced from the input data rather than from the output of the first convolution layer.

**Fig 1.**
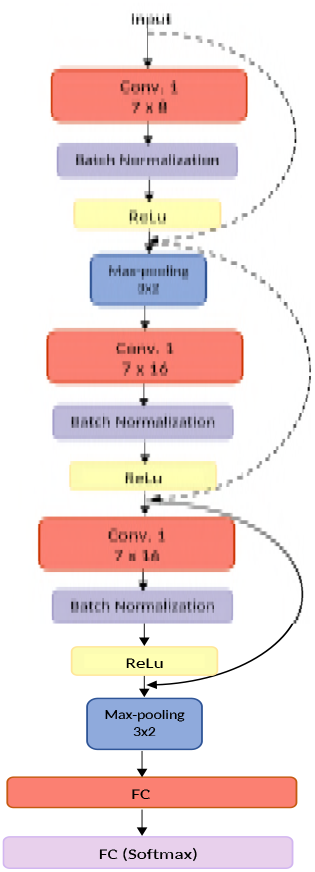
S-ResNet architecture, 3 convolutions (Conv), 2 Fully Connected (FC), and 3 Shortcuts. The numbers inside convolution layer boxes show the filter size x number of filters and in the max-pooling layer filter size x stride. We followed the same notation as in the original ResNet work for the shortcut by using dotted lines for projection shortcut and solid line for identity shortcut.

### Comparison of S-ResNet with Traditional Machine Learning Methods

Classical machine learning methods such as Logistic Regression (LG) [16], Support Vector Machine (SVM) [16], [17], [15] and Random Forests (RF) [18] [19] have shown good performance for splice site recognition. Among deep learning models we have implemented (see table 1), we selected S-ResNet to benchmark with classical methods since it showed the best performance as we will show below. The hyper-parameters were extensively tuned for the three baseline methods using the training and a validation set to reach the best result. Seven subsets have been selected randomly with the following sizes: 1,000; 3,000; 10,000; 30,000; 100,000; 200,000 and 300,000 samples. For an accurate comparison, the exact same training and test datasets have been used to evaluate all techniques. Six evaluation metrics have been measured: accuracy, training time, area under curve (AUC), Area Over Precision Recall Curve (aoPRC), sensitivity and specificity.

## Results

### S-ResNet vs. Traditional Classifiers

Traditional baseline machine learning techniques do not intrinsically handle the sequential nature of the DNA data. In addition, they have limited scalability when the number of examples becomes very large. Despite of that, they showed good result in this evaluation. Figure 2 compares the result of S-ResNet and the three baselines on test dataset for AUC and additional results for other metrics are provided in the supplementary S.1. It is out of the scope of this work to include techniques which incorporate the sequence information in classical machine learning methods such as k-mer SVM etc. Our purpose is to use directly raw sequence data and show the benefit of deep learning compared with manual feature engineering. S-ResNet has led to the best performance for most of the evaluated metrics starting from 30,000 examples and achieving 97.52 AUC for largest dataset of size 300,000, this is 2% better than LR the second best result. Additionally, S-ResNet has an average of accuracy 92.5% (for 100,000) and 93% (for 200,000 and 300,000) resulting in about 2% and 3% improvement compared to second best result obtained by SVM and LR. Although SVM and LR have almost the same result in the accuracy staring from 10,000 samples, there is a large difference between them in terms of AUC. However, when relatively small data size (10,000 samples and less) is used, S-ResNet ranked the second or the third best technique following LR and SVM.

**Fig 2.**
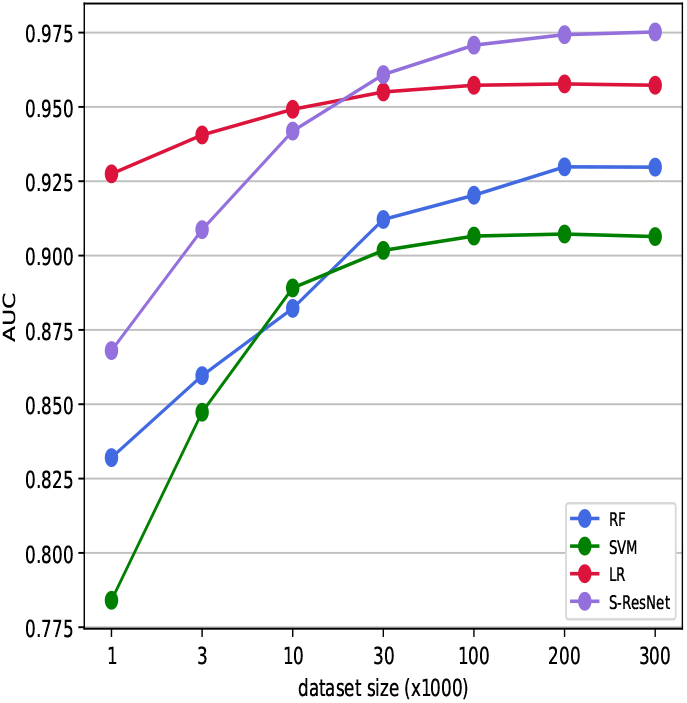
Comparing (in terms of AUC) of traditional classifiers, i.e., RF, SVM, LR with the proposed S-ResNet model.

### Role of Model Depth

In this Analysis, we tested different architecture depths. For the evaluation of performance we used the following metrics: Accuracy, AUC, number of trainable parameters, number of floating point operations, specificity, and sensitivity. These metrics were calculated based on test dataset. In each experiment the data is split randomly into three subsets: training, validation and test sets with the following proportions: 40%-20%-40% respectively. In the first set of experiments, the number of training epochs fixed to 300. The evaluation is repeated 5 times for different random splits of the dataset, this allowed to estimate the average and standard deviation of the different metrics. The result summary is given in the Table S.1 in supplementary material. Shallow architecture of each model family has shown better performance compared to deep architecture. ResNet-18 and ShallowNet gave the highest AUC and accuracy, while VGG models had the worst AUC. DeepBind and AlexNet models ranked in between. In terms of model complexity (number of trainable parameters and floating point operations), shallow models, i.e., DeepBind, ShallowNet and S-ResNet have the least computational complexity.

Because the best training convergence differs from one architecture to another and not necessary to achieve the best performance at 300 epochs, in the second set of experiments, early stopping is used with patience of 10 epochs. Loss on validation set is used to decide on stopping the training if no improvement is noticed. The metrics average and standard deviation were based on 10 repetitions of the experiments and the results are presented in Table 2. Early stopping improved the performance (accuracy and AUC) for all the models. It is known that early stopping helps preventing from over-fitting. Overall S-ResNet had the highest performance with AUC (97.6) and accuracy (93.4). VGGNet-E improved significantly from 94.1 AUC with 300 epochs to 97.4 AUC with early stopping to rank the second with ShallowNet, ResNet-18 and ResNet-34.

**Table 2.** Comparison of models performance for metrics: Accuracy (Acc.), Area Under ROC curve (AUC), number of floating point operations (# of FLOPS), number of trainable model parameters (# of parameters) during training early stopping is used with patience of 10 epochs. The metrics are estimated based on the test dataset averaged across 10 repetitions.

| Model | Acc. | AUC | # of FLOPS | # of parameters | Sensitivity | Specificity |
| --- | --- | --- | --- | --- | --- | --- |
| ShallowNet | 93.1 ± 0.2 | 97.4 ± 0.04 | **6,399** | **3,202** | **0.929 ± 0.007** | 0.932±0.01 |
| S-ResNet | **93.4 ± 0.1** | **97.6 ± 0.04** | 11,060 | 5,324 | **0.929 ± 0.007** | 0.938±0.01 |
| ResNet-18 | 92.6 ± 0.8 | 97.4±0.2 | 1,219,450 | 243,986 | 0.896±0.03 | 0.956±0.01 |
| ResNet-34 | 91.8±1.6 | 97.4±0.2 | 2,288,330 | 457,202 | 0.873±0.05 | **0.963 ± 0.01** |
| AlexNet-5 | 92.4±0.6 | 96.9±0.4 | 26,471 | 13,238 | 0.923±0.02 | 0.925±0.02 |
| AlexNet-6 | 92.6±0.1 | 97.1 ± 0.1 | 49,767 | 24,950 | 0.911±0.01 | 0.940±0.01 |
| VGGNet-A | 92.7±0.4 | 96.9 ± 0.5 | 207,543 | 104,262 | 0.906±0.03 | 0.947±0.02 |
| VGGNet-E | 93.1±0.3 | 97.4 ± 0.1 | 467,511 | 234,718 | 0.917±0.02 | 0.945±0.02 |
| DeepBind | 92.4±0.3 | 96.9 ± 0.08 | 3,655 | 1,854 | 0.927±0.01 | 0.920±0.01 |

### Result Summary

Figure 3 provides performance summary of AUC (y-axis), computational complexity (FLOPS) (x-axis) and number of parameters (diameter of the circle). Figure 4 shows the ROC with early stopping. The results were very close to each other with outperforming for S-ResNet and the lowest performance for DeepBind. We captured the cost and the accuracy at each epoch during model training. We also computed the validation and test costs as well as accuracies during training at each epoch. This allowed us to monitor the model behaviour during training. Figure 5 and Figure 6 show the cost and accuracy with early stopping for the different models. The accuracy and cost for validation and test datasets have similar trend, however, the models do not have the same curve length due to the use of early stopping.

**Fig 3.**
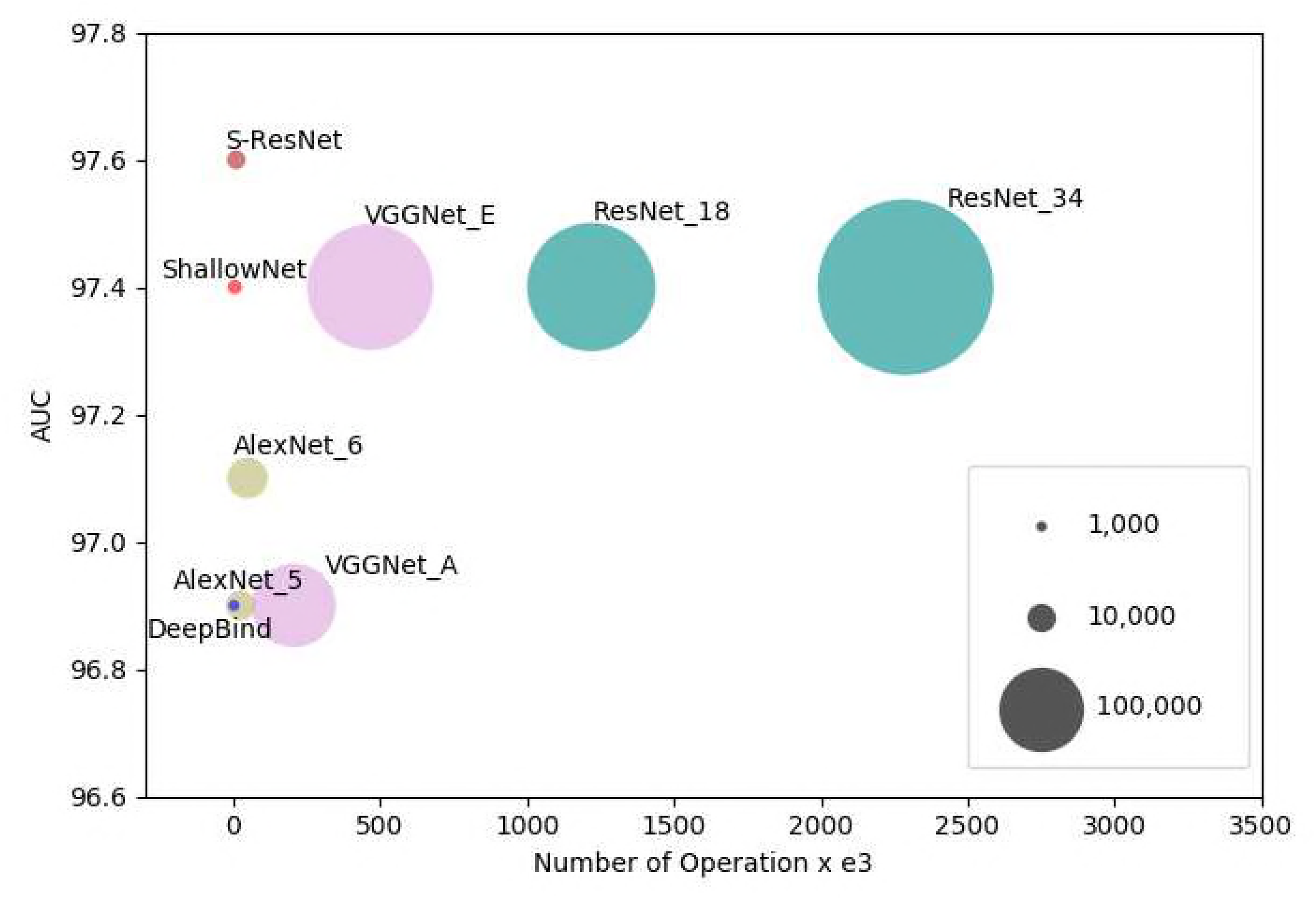
Comparison of model performance using three metrics: Number of floating point operations (X_axis), AUC score (Y_axis) and number of tunable model parameters (circle diameter).

**Fig 4.**
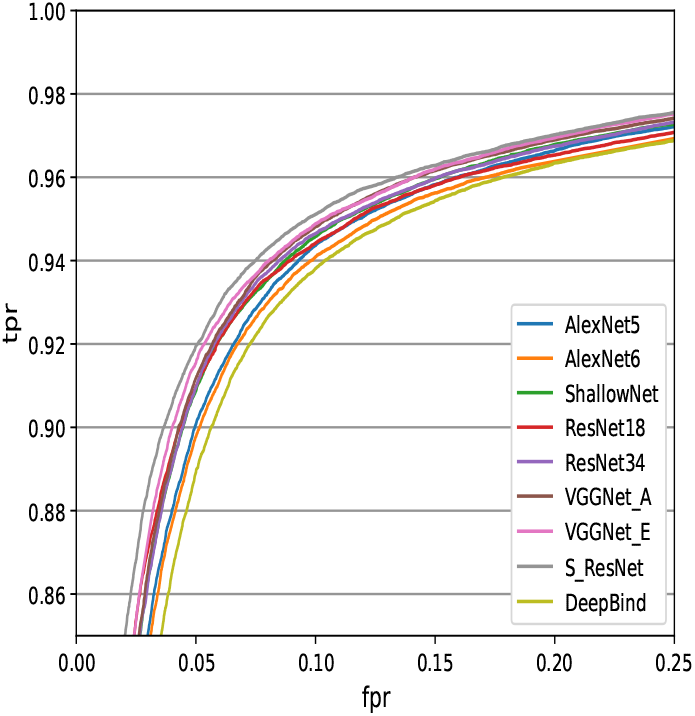
Average ROC curves estimated on the test sets using early stopping function for the training. The scale is zoomed to show the difference between the evaluated models.

**Fig 5.**
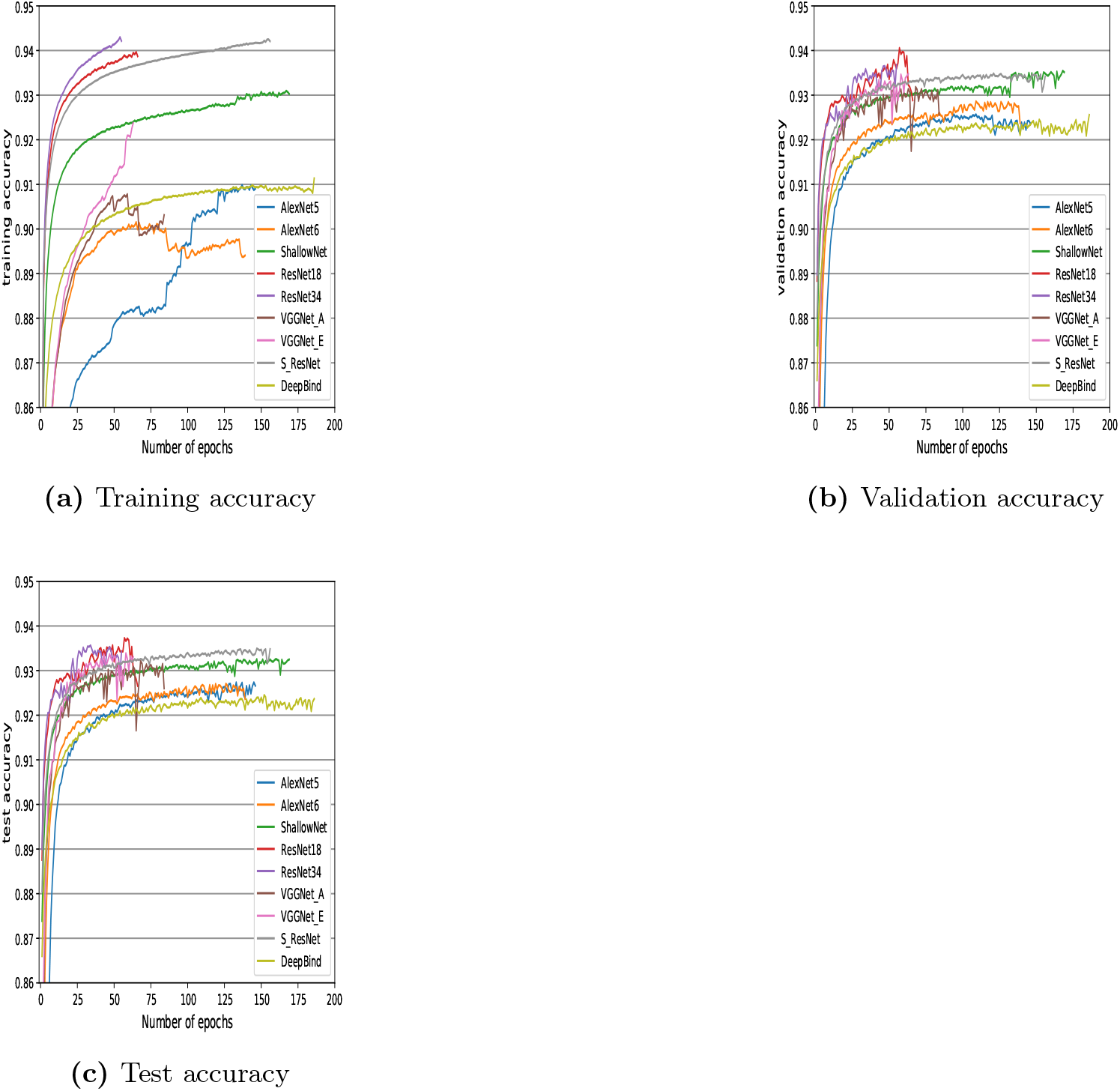
Training, validation and test accuracies during model training with early stopping.

**Fig 6.**
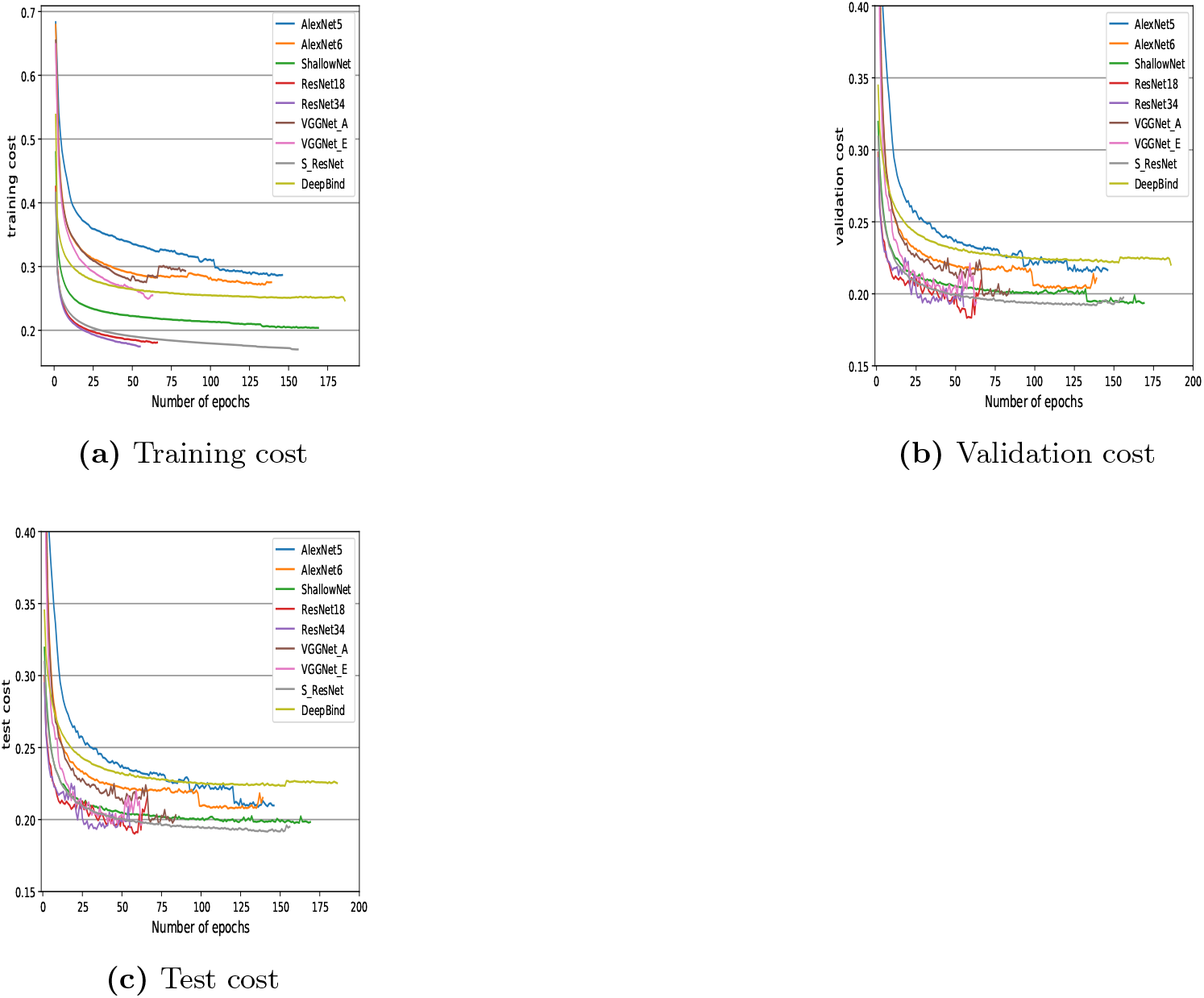
Training, validation and test costs during model training with early stopping.

## Discussion

In this paper, we adapted known deep learning models from computer vision for the classification of splice sites based on raw DNA sequence. Our analysis investigated the role of model depth for this classification task. We proposed a new model named S-ResNet inspired by shallow ResNet. We provided comparison of S-ResNet with traditional machine techniques Logistic Regression (LR), Support Vector Machines (SVM) and Random Forests (RF). S-ResNet gave the best performance for large dataset (30,000 samples and larger) while LG and SVM gave the best results for small dataset.

### Role of Model Depth

Our experiments showed that increasing architecture depth, for the three studied deep learning model families (AlexNet, VGG and ResNet), did not improve the performance of splice site classification based on raw DNA sequences. This result might appear contradicting with the known importance for depth in deep learning literature. However the results shown in the literature tackled different classification problem where one of the differences compared to this work lays in the number of classes in the dataset. Typically in computer vision, speech or text dataset the number of classes is large and vary between 10s to 100s or more of classes, whilst, the splice site dataset used in this paper has two classes (spice and non-splice DNA sequences). We plan to further explore this justification in future work by considering other genomic datasets with large number of classes. Overall, the proposed S-ResNet model gave the best performance. The shortcut connection present in S-ResNet architecture combined with it shallow architecture seem to contribute in solving the gradient vanishing and accuracy degradation problems seen in deep architectures. S-ResNet showed performance improvement compared with other shallow networks such as ShallowNet. VGGNet-E showed close performance to S-ResNet and equivalent to ShallowNet but exhibited high computational cost due to its increased number of parameters and operations.

Figures 5 (b,c) and 6 (b,c) show superiority of ResNet-18 and ResNet-34 over S-ResNet and ShallowNet. However, on average the accuracy is higher for S-ResNet and ShallowNet because of longer training needed (larger epochs) with a slight increase in accuracy and cost. The original work for ResNet [3] with an imaging dataset, ResNet-34 has smaller training and validation error than ResNet-18. This result is also valid in our experiment for the training cost (Figure 6 (a)), and not all the time for validation and test result were the two curves keep fluctuating (Figure 6 (b,c)).

### Over-fitting

The similarity in training and validation curves (or the improving with the validation data with some cases) indicates that over-fitting is well controlled in our analysis. Dropout function was used as a regularizer with all the experiment except S-ResNet. We did more investigation on S-ResNet and test its performance with dropout regularizer. Figure 7a and 7b compare the cost and accuracy for the validation data with and without dropout for S-ResNet. With dropout, the AUC and accuracy were 97.5 and 93.5 while without Dropout were 97.6 and 93.4 respectively. We notice that the required number of epochs for training to stabilize increased when Dropout is used from 133 (without Dropout) to 148 (with Dropout) and convergence time increased from 89 min (without Dropout) to 119 min (with Dropout).

**Fig 7.**
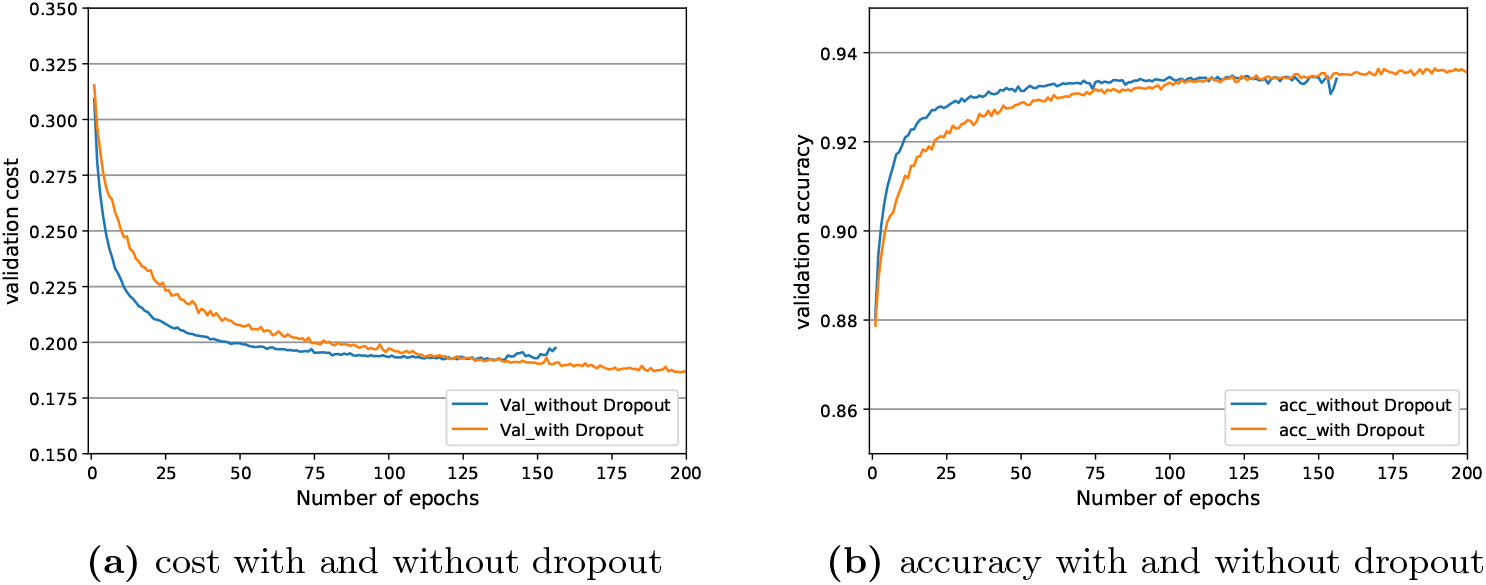
Comparison of accuracy and cost with/with-out dropout

### Speed of Convergence

We did not include computational time in our list of performance metrics because we ran the experiments on systems with different GPU specifications. The number of floating point operations and length of training (number of epochs with early stopping training) can be used as a proxy to estimate computational time (Table 2 and table 3). We note although ResNet and VGGNet required the least epochs in training they are more computationally expensive compared to shallow architectures due to the large number of parameters and operations involved in the model. In general, shallow architectures required more epochs for training than deep models.

**Table 3.** Number of epochs needed for each model for early stopping.

| Model | # of epochs |
| --- | --- |
| ShallowNet | 170 |
| S-ResNet | 132 |
| ResNet-18 | 38 |
| ResNet-34 | 33 |
| AlexNet-5 | 93 |
| AlexNet-6 | 93 |
| VGGNet-A | 55 |
| VGGNet-E | 49 |
| DeepBind | 146 |

### Effect of Long Model Training in ResNet

ResNet is known for the nice property of continuous validation loss reduction when trained with few hundreds epochs. We conducted an additional experiment to evaluate the behavior of ResNet model during training for 1000 epochs. The results of this experiment are summarized in Figure ?? for accuracy and cost. The validation and test curves are overlapping for ResNet-18 and ResNet-34 for both accuracy and cost figures. This result is expected as no early stopping or parameter tuning was used on validation. In this case validation and test sets are equivalent. For ResNet-18, validation cost and accuracy improved rapidly until 200 epochs before a slow training phase was then observed. No improvement could be seen beyond 400 epochs and the convergence is rather noisy in this phase. The training and validation curve are close to each other indicating that no over-fitting is observed. For ResNet-34, a faster convergence is noted in the early rounds of training. This is explained by the higher model capacity due its extended trainable parameters. However validation cost and accuracy degrades dramatically beyond 50 epochs. The model is clearly over-fitting the training data and further training the model is only increasing the gap between training and validation cost. This experiment showed that deep ResNet (ResNet-34) is not best suited for this classification task. Also lengthy training of ResNet worked moderately well for ResNet-18. Early stopping showed overall better result with ResNet as it prevented over-fitting.

## Conclusion

Inspired by the advances of deep learning in computer vision, we adapted successful models, i.e., AlexNet, ResNet and VGG, for the classification of splice sites from raw DNA sequence. For each model family, we defined three architecture depths based on the number of convolution layers used. We introduced a shallow version of ResNet, called S-ResNet suited for our classification task. S-ResNet has both advantage of shallow architecture and shortcut connection. Shallow models have shown an overall better performance than models with deep architecture. S-ResNet ranked top and had slightly better performance than the second model (ShallowNet). We conclude that splice site classification does not require deep architecture and increasing the number of convolution layers merely increases the computational cost without performance gain. As future work, we plan to further investigate the role of depth in other classification tasks using genomic sequence data. We will consider problems with larger number of classes. We will also explore other models based on Recurrent Neural Networks.

## Supporting Information

**Fig S.1.**
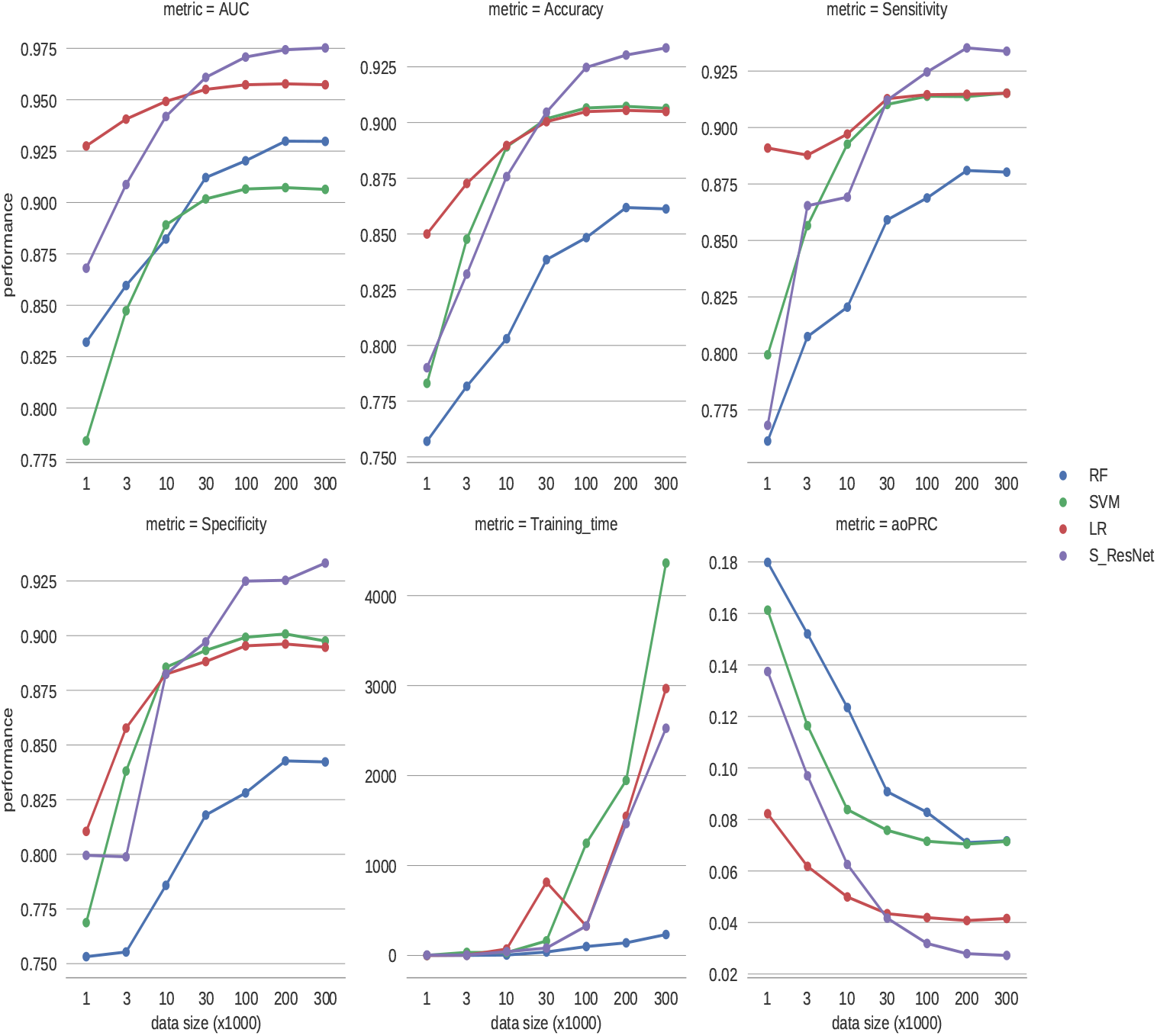
Comparison of S-ResNet with traditional machine learning techniques.

**Table S.1.**
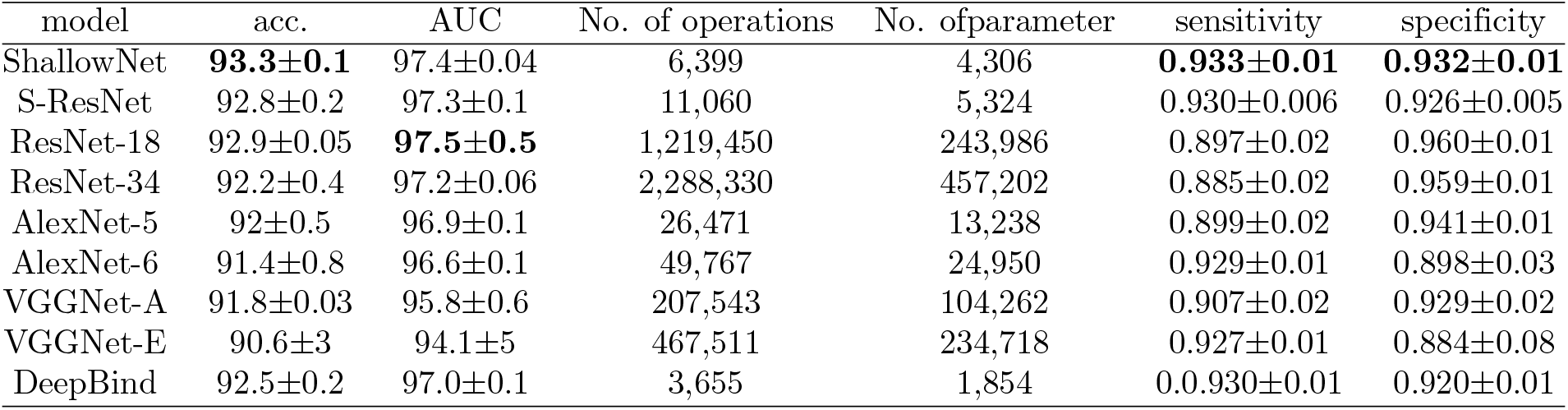
Shallow and deep Models metrics comparison with training for 300 epochs.

| model | acc. | AUC | No. of operations | No. of parameter | sensitivity | specificity |
| --- | --- | --- | --- | --- | --- | --- |
| ShallowNet | **93.3±0.1** | 97.4±0.04 | 6,399 | 4,306 | **0.933±0.01** | **0.932±0.01** |
| S-ResNet | 92.8±0.2 | 97.3±0.1 | 11,060 | 5,324 | 0.930±0.006 | 0.926±0.005 |
| ResNet-18 | 92.9±0.05 | **97.5±0.5** | 1,219,450 | 243,986 | 0.897±0.02 | 0.960±0.01 |
| ResNet-34 | 92.2±0.4 | 97.2±0.06 | 2,288,330 | 457,202 | 0.885±0.02 | 0.959±0.01 |
| AlexNet-5 | 92±0.5 | 96.9±0.1 | 26,471 | 13,238 | 0.899±0.02 | 0.941±0.01 |
| AlexNet-6 | 91.4±0.8 | 96.6±0.1 | 49,767 | 24,950 | 0.929±0.01 | 0.898±0.03 |
| VGGNet-A | 91.8±0.03 | 95.8±0.6 | 207,543 | 104,262 | 0.907±0.02 | 0.929±0.02 |
| VGGNet-E | 90.6±3 | 94.1±5 | 467,511 | 234,718 | 0.927±0.01 | 0.884±0.08 |
| DeepBind | 92.5±0.2 | 97.0±0.1 | 3,655 | 1,854 | 0.930±0.01 | 0.920±0.01 |

**Fig S.2.**
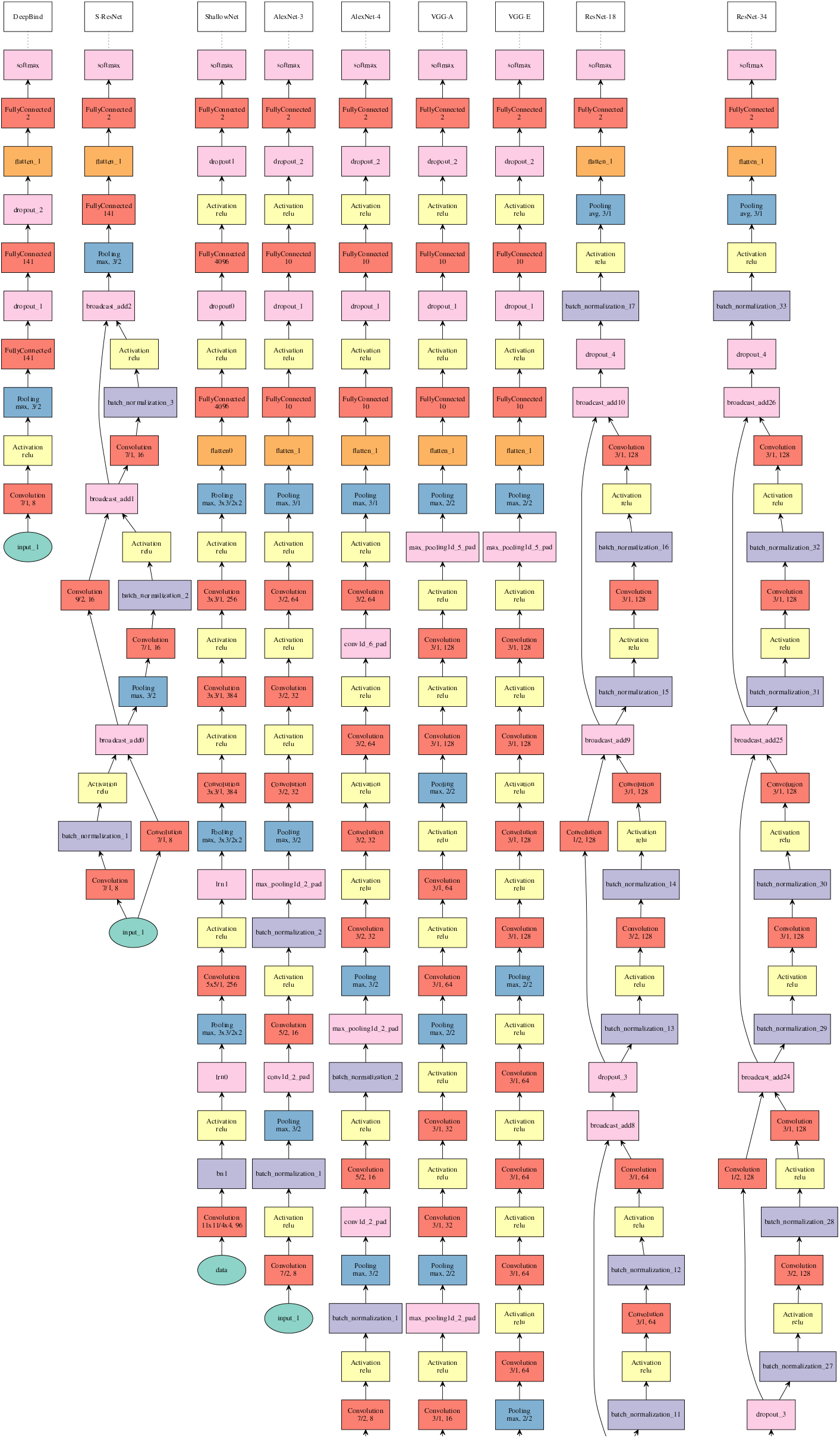
Model Architectures

**Fig S.3.**
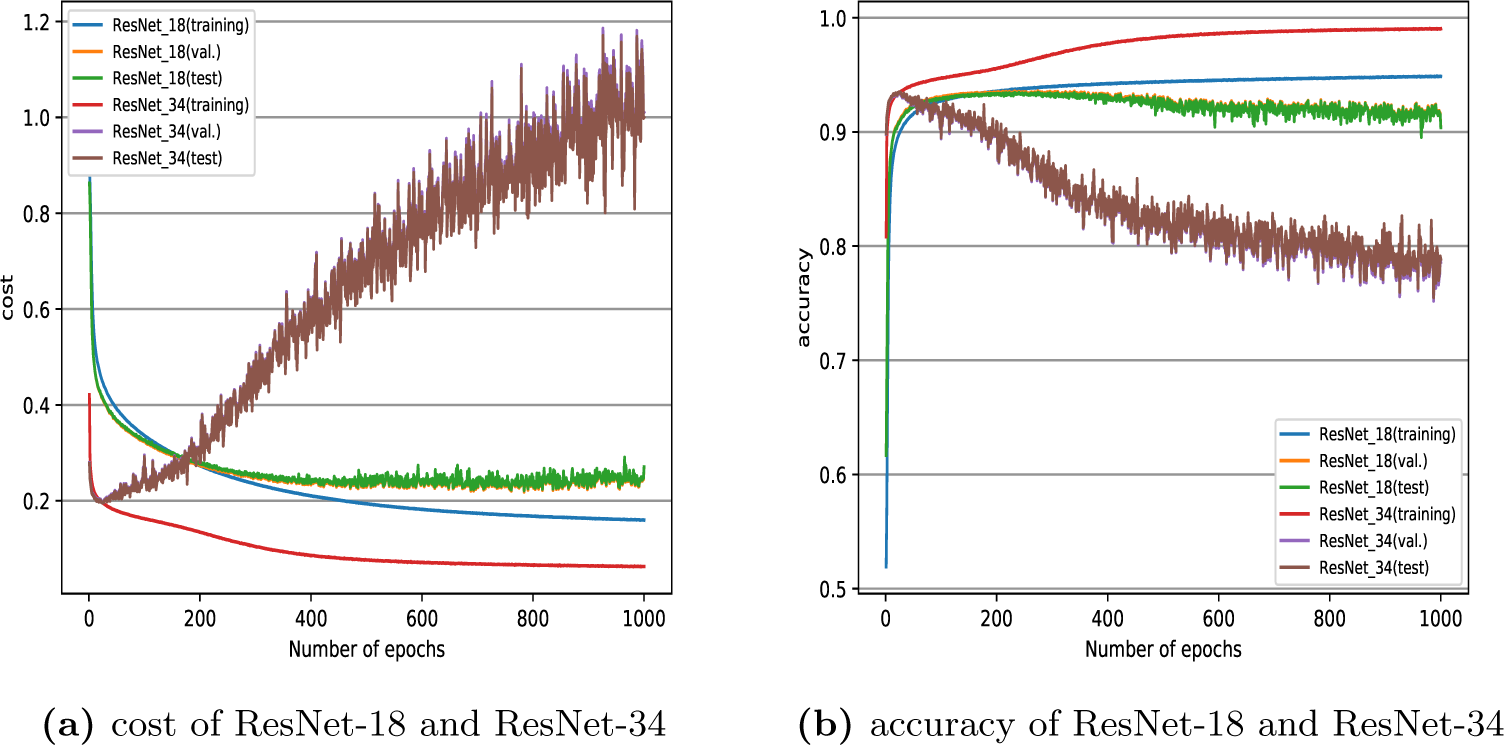
Summary of accuracy and cost of ResNet-18 and ResNet-34 for training with 1000 epochs

